# Ethylene signaling increases reactive oxygen species accumulation to drive root hair initiation in Arabidopsis

**DOI:** 10.1101/2021.11.14.468514

**Authors:** R. Emily Martin, Eliana Marzol, Jose M. Estevez, Gloria K. Muday

**Affiliations:** Departments of Biology and Biochemistry and the Center for Molecular Signaling, Wake Forest University, 1834 Wake Forest Road, Winston-Salem, NC 27109; Fundación Instituto Leloir and IIBBA-CONICET. Av. Patricias Argentinas 435, Buenos Aires C1405BWE, Argentina; ANID - Millennium Science Initiative Program - Millennium Institute for Integrative Biology (iBio) and Millennium Nucleus for the Development of Super Adaptable Plants (MN-SAP), Santiago, Chile

**Keywords:** Ethylene, reactive oxygen species, root hairs, peroxidase, RBOH

## Abstract

Root hair initiation is a highly regulated aspect of root development. The plant hormone, ethylene, and its precursor, 1-amino-cyclopropane-1-carboxylic acid (ACC), induce formation and elongation of root hairs. Using confocal microscopy paired with redox biosensors and dyes, we demonstrated that treatments that elevate ethylene levels led to increased hydrogen peroxide accumulation in hair cells prior to root hair formation. In two ethylene-insensitive mutants, *etr1-3* and *ein3/eil1*, there was no increase in root hair number or ROS accumulation. Conversely, *etr1-7*, a constitutive ethylene signaling receptor mutant, has increased root hair formation and ROS accumulation like ethylene-treated Col-0 seedlings. The *caprice* and *werewolf* transcription factor mutants have decreased and elevated ROS levels, which are correlated with levels of root hair initiation. The *rhd2-6* mutant, with a defect in the gene encoding a ROS synthesizing Respiratory Burst Oxidase Homolog C (RBOHC) and the *prx44-2* mutant defective in a class III peroxidase, showed impaired ethylene-dependent ROS synthesis and root hair formation and EIN3/EIL1 dependent transcriptional regulation. Together, these results indicate that ethylene increases ROS accumulation through RBOHC and PRX44 to drive root hair formation.

**SUMMARY STATEMENT:** The gaseous hormone ethylene increases root hair initiation by elevating reactive oxygen species (ROS) in trichoblast cells. Genetic and biochemical approaches identified ethylene-regulated ROS producing enzymes that drive root hair initiation.

## INTRODUCTION

The initiation of root hairs is genetically programmed and environmentally sensitive, making them an ideal model for studying single cell differentiation in plants. Root hairs are single-cell extensions that differentiate from longitudinal epidermal cell files, known as trichoblasts (Leavitt, 1904; Salazar-Henao et al., 2016). The formation of root hairs is modulated by environmental changes to increase root surface area to allow for efficient water and nutrient uptake (Bruex et al., 2012), while also anchoring plants in soil to reduce erosion (De Baets et al., 2020). In Arabidopsis, the root epidermis consists of an alternating pattern of trichoblasts, which form root hairs, and atrichoblasts, which are non-hair forming cells (Pemberton et al., 2001; Salazar-Henao et al., 2016). Root hair formation is dictated by cell positioning; epidermal cells overlying two cortical cells can become hair forming cells, while those overlying one cortical cell will become non-hair cells (Berger et al., 1998; Salazar-Henao et al., 2016). Root hair development is separated into two processes: root hair initiation and root hair elongation (Dolan et al., 1994). A recent report divided this process into 10 precise stages and examined the molecular mechanisms that drive this process. During stages −7 to −1, RHO Of Plants (ROP) proteins (ROP2, ROP4, and ROP6) and ROP Guanine Nucleotide Exchange factors (ROP GEFs) that regulate ROP activity accumulate at the future site of initiation (Denninger et al., 2019; Molendijk et al., 2001), this is followed by polymerization of actin filaments along which vesicles move to deposit membrane needed for polarized tip growth (Salazar-Henao et al., 2016). During tip growth, tip focused reactive oxygen species (ROS) (Foreman et al., 2003; Gayomba and Muday, 2020; Monshausen et al., 2007) and Ca^2+^ gradients (Carroll et al., 1998) have been identified to drive exocytosis of the cell wall and membrane materials driving subsequent root hair elongation. These processes are separable as mutants with either impaired root hair initiation or elongation have been identified (Masucci and Schiefelbein, 1994; Schiefelbein and Somerville, 1990).

Genetic screens in *Arabidopsis thaliana* have provided a wealth of insight into the proteins that drive root hair development (Lee and Schiefelbein, 1999; Masucci and Schiefelbein, 1996; Rerie et al., 1994). For example, the mutants *transparent testa glabra (ttg), glabra2 (gl2)*, and *werewolf (wer)* have root hairs that form from both trichoblast and atrichoblast cells (Di Cristina et al., 1996; Galway et al., 1994; Lee and Schiefelbein, 1999). Many of the protein products of these mutants have been mapped to transcriptional cascades that drive root hair differentiation (Shibata and Sugimoto, 2019) and have been shown to localize prior to root hair initiation (Denninger et al., 2019). In non-hair cells, a transcriptional complex comprised of three transcription factors (TFs), WER, GLABRA 3 (GL3) or its functionally redundant ENHANCER OF GLABRA (EGL3), and TTG1, function as a transcriptional activator of the GL2 protein, which leads to repression of root hair initiation (Grebe, 2012; Salazar-Henao et al., 2016). In trichoblasts, *WER* expression is repressed, which allows for the formation of an alternate transcriptional complex comprised of CAPRICE (CPC) or the functionally redundant proteins ENHANCER OF TRY AND CPC1 (ETC1), ETC3, or TRYPTICHON (TRY) (Schiefelbein et al., 2014). When this pathway is active, GL2 is not expressed and root hair initiation proceeds (Salazar-Henao et al., 2016).

Another genetic screen identified the *root hair defective* (*rhd*) mutants which have impaired root hair initiation, elongation, or structure (Masucci and Schiefelbein, 1994; Schiefelbein and Somerville, 1990). The mutation in *rhd2*, was mapped to the *RBOHC* (respiratory burst oxidase homolog C/NADPH oxidase) gene (Foreman et al., 2003). The *rhd2* mutant was identified for altered root hair elongation (Schiefelbein and Somerville, 1990), but was recently reported to also have impaired root hair initiation (Gayomba and Muday, 2020). RBOHs are integral plasma membrane proteins that produce superoxide, which can be dismutated to hydrogen peroxide via superoxide dismutase (SOD) or other non-enzymatic mechanisms (Chapman et al., 2019). H_2_O_2_ can then enter into the cell through aquaporins (Bienert et al., 2007), where it can act as a signaling molecule to drive cellular processes. Signaling induced ROS regulates protein function by reversibly oxidizing cysteine residues to sulfenic acids (Cys-SOH) (Poole and Schoneich, 2015).

There are 10 RBOH family members (RBOHA-RBOHJ) in Arabidopsis and each play distinct roles in organ development and stress response (Chapman et al., 2019). RBOHs can be regulated transcriptionally or enzymatically by a number of mechanisms, including calcium binding, phosphorylation, and phosphatidic acid binding (Kobayashi et al., 2007; Postiglione and Muday, 2020; Suzuki et al., 2011). RBOH-induced ROS production is also regulated by hormone signaling, as many plant hormones generate ROS as a mechanism to drive growth and developmental processes (Chapman et al., 2019; Kwak et al., 2003; Mittler et al., 2011; Postiglione and Muday, 2020); For example, abscisic acid (ABA), a hormone involved in abiotic stress response, has been shown to induce RBOH-derived ROS production to prevent water loss in leaves (Kwak et al., 2003). Auxin induces root hair initiation (Gayomba and Muday, 2020) and elongation (Mangano et al., 2017) through a localized increase in ROS, suggesting that RBOH may drive hormone induced root hair elongation.

The plant hormone ethylene enhances root hair initiation and elongation. Treatment with ethylene, or its precursor 1-amino-cyclopropane-1-carboxylic acid (ACC), leads to proliferation of root hairs, with substantial increases in their length (Tanimoto et al., 1995). The ethylene induction of root hairs occurs through the canonical ethylene signaling pathway, which is initiated when ethylene binds to one of the five receptors, ETR1, ERS1, ETR2, ERS2, or EIN4 (Binder, 2020; Bleecker, 1999). When ethylene is absent, the receptors are in the “on state” leading to activation of the CTR1 Raf-like kinase (Kieber et al., 1993); this turns the pathway off via phosphorylation and subsequent degradation of the EIN2 transmembrane protein (Alonso et al., 1999; Ju et al., 2012). When ethylene binds to its receptors they shift to the “off state” which prevents activation of CTR1. This allows for cleavage and translocation of the EIN2 C-terminus into the nucleus (Ju et al., 2012; Wen et al., 2012) where it stabilizes the EIN3/EIL1 TFs leading to ethylene responsive gene expression. Previous work has shown that ethylene regulates root hair elongation through the EIN3 transcription factor (Feng et al., 2017) and that EIN3 also controls the formation of ectopic root hairs when ethylene reaches very high levels (Qiu et al., 2021). EIN3 physically interacts with RHD6, a positive regulator of root hair development, to form a transcriptional complex that binds to and induces expression of *RSL4* resulting in increased root hair elongation (Feng et al., 2017). However, the mechanistic events that drive ethylene-induced root hair initiation have not been fully described and the role of ROS as a downstream molecule in ethylene-induced root hair development has not been reported.

This work asked whether ethylene acts to increase ROS levels to drive root hair initiation. Fluorescent dyes and biosensors that report ROS levels were used to examine ROS accumulation after ethylene or ACC treatment in trichoblast cells in the differentiation zone. This response was shown to be dependent on ETR1 receptor activity and EIN3 TF activity. The role of RBOHC and class III peroxidase enzymes in ethylene-dependent ROS synthesis was demonstrated using mutants in genes encoding these ROS generating enzymes. Together these experiments demonstrated that ROS is a signaling molecule in ethylene induced root hair formation and identified several enzymes that participate in producing ethylene-induced ROS.

## RESULTS

### Root hair numbers increases in ACC treated roots

The role of ethylene signaling in root hair initiation was examined by testing the effects of short-term treatments with the ethylene precursor, ACC, on the number and position of root hairs in 5-day old seedlings. Root hairs were visualized in wild type (Col-0) seedlings grown in the presence of 0.7 µM ACC for 4 hours (Fig. 1A). The root tip was divided into three 500 µm zones starting from the root tip and the number of initiated root hairs formed in each zone was quantified Fig. 1B). We defined root hair initiation as root hairs that were at stage +2 and above as described previously (Denninger et al., 2019). In zone 1, root hairs did not form in either untreated or ACC-treated seedlings. In zone 2, there were very few root hairs in untreated roots, however, ACC-treatment increased the number of root hairs by 10-fold. There was also a 2-fold increase in root hair number in zone 3 of ACC-treated seedlings compared to untreated controls. This dose of ACC does not cause ectopic root hair formation in non-hair cells, but rather increased the number of hair cells forming root hairs in these two zones. It is also evident from Fig. 1, that this treatment increased the length of root hairs in both zone 2 and 3, consistent with prior reports (Feng et al., 2017; Harkey et al., 2018). These data suggest that ACC-induced root hair initiation begins between 500 and 1000 µM from the root tip. Therefore, to understand the mechanisms driving the process of root hair initiation our experiments focused on this region.

**Figure 1.**
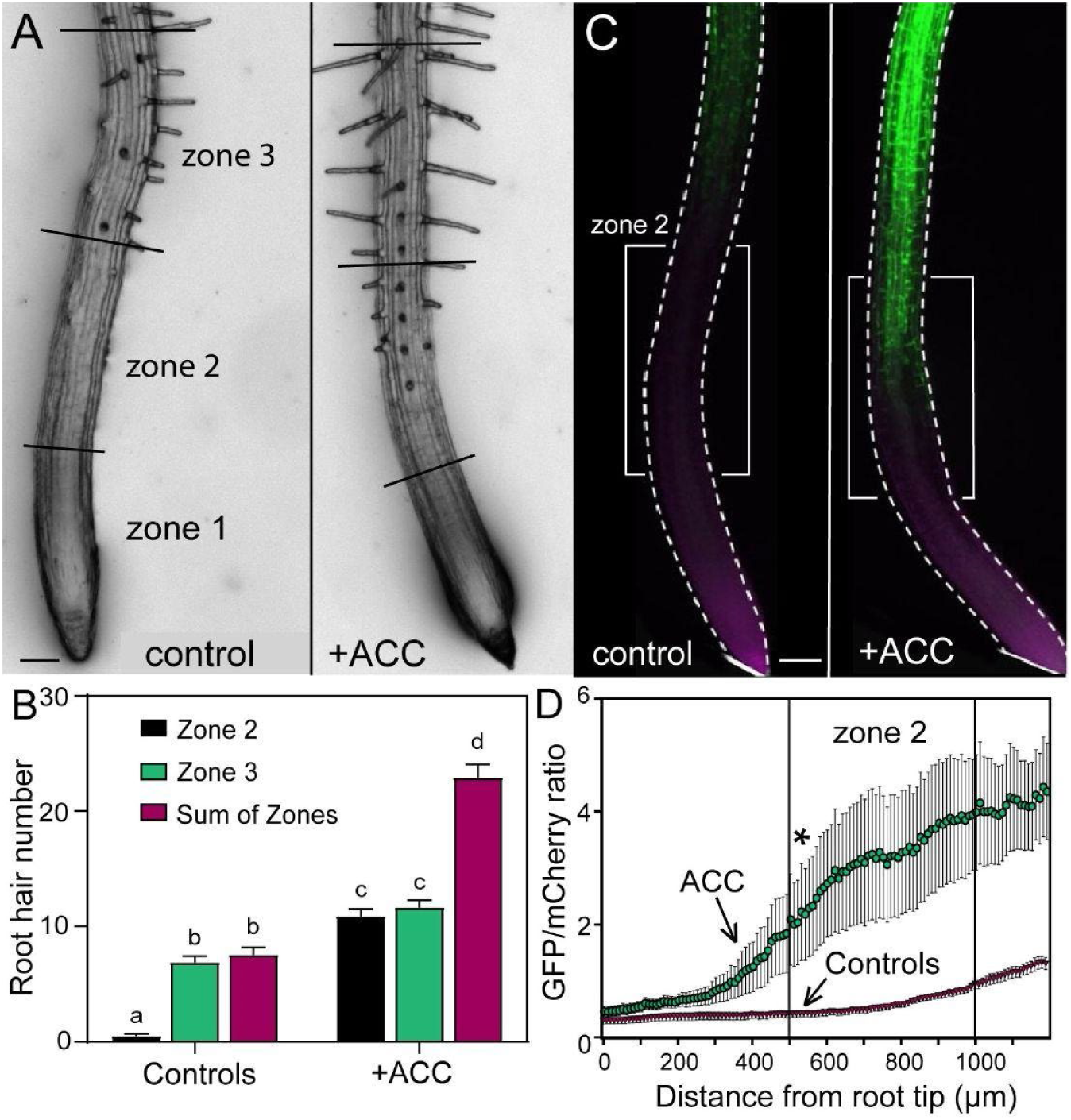
ACC treatment increases root hair number and ROS dependent gene expression along the root after 4 hours. (A) Representative images of root tips of Col-0 treated with and without ACC for 4 hours. Zones 1, 2, and 3 represent 500 µm root sections. Scale bar for all images are 100 µm. (B) Root hair quantification of untreated and 4h ACC treated seedlings. No root hairs formed in zone 1. Data are means ± SEM of total RH in zone 2, zone 3, and all zones from 3 experiments (n=18-24 seedlings per experiment). Columns with different letters indicate statistically significant differences determined by two-way ANOVA followed by Tukey’s multiple comparisons test. (C) Representative images of root tips of Col-0 containing the ZAT12p-ROS reporter treated with or without ACC for 4 hours with GFP signal in green and mCherry signal in magenta. White brackets indicate zone 2 of the root. (D) Quantification of the ratio of GFP/mCherry fluorescence intensity using a line profile along the roots of an average of 6-12 seedlings of 3 independent experiments of untreated or ACC treated seedlings. Data are means ± SEM. The asterisk represents the shortest distance from the root tip at which the GFP fluorescence in the presence of ACC values became significantly different from the controls as determined by student’s t-test (at approximately 510 µm).

Long term ACC and ethylene treatment also result in a shorter primary root due to reduced elongation of root cells, so we examined whether the effect on root hair initiation was the result of altered length (Harkey et al., 2018). The effect of 4 hours of treatment with low doses of ACC (0.7 µM) on root elongation were examined. Primary roots were stained with the cell wall specific dye, propidium iodide (PI). We then measured the length of 5 epidermal cells from 6-8 roots per treatment condition, measuring the length of cells at either end and in the middle of zone 2 (designated zone 2A, 2B, and 2C in Fig. S1). There was no difference in cell length in any of these 3 regions after 4 hours of ACC treatment compared to untreated controls. These results are consistent with short term and low dose ACC treatments increasing root hair number by inducing root hair formation from trichoblasts in zone 2, rather than as an indirect effect of a shorter primary root.

### A ROS dependent transcriptional reporter increases in ACC treated roots

We asked whether ethylene leads to elevated reactive oxygen species (ROS) to drive ethylene-induced root hair initiation. ROS dependent gene expression along the root was examined using the ZAT12p-ROS ratiometric biosensor (Lim et al., 2019). This reporter construct contains the promoter of the ROS sensitive transcription factor, *ZAT12*, driving GFP and the constitutive ubiquitin10 promoter driving mCherry. We compared the fluorescent signal of *ZAT12p*-GFP in the presence and absence of ACC as visualized by laser scanning confocal microscopy (LSCM), with GFP reported as green and mCherry reported as magenta (Fig. 1C). The ratio of signal of GFP (green) to mCherry was quantified across the entire root as a distance from the root tip. In ACC-treated roots, the GFP/mCherry ratio increased beginning at approximately 500 μm from the root tip, which corresponded to zone 2, where root hair induction was maximal upon treatment with ACC (Fig. 1D). To examine the possibility that the *ZAT12* promoter is ethylene regulated (rather than being ROS regulated), we examined transcriptomic datasets in which roots were treated with ACC or ethylene (Harkey et al., 2018). These three data sets do not show ACC or ethylene-driven transcriptional changes in *ZAT12* transcript abundance.

### Ethylene increases the signal of the H_2_O_2_ selective dye, Peroxy Orange1, in trichoblast cells

The hydrogen peroxide selective dye, Peroxy-Orange 1 (PO1), was used to determine whether there were cell type-specific increases in ROS in response to treatment with ACC or ethylene gas. PO1 is a permeable, boronate-based dye that is non-fluorescent in its reduced form, but becomes fluorescent when irreversibly oxidized by H_2_O_2_ (Dickinson et al., 2010). Col-0 seedlings were treated with either 0.7 µM ACC or 0.05 ppm of ethylene gas for 4 hours followed by PO1 staining. In Col-0 roots, PO1 fluorescence was visualized using LSCM in zone 2 of roots treated with control, ACC, or ethylene gas for 4 hours (Fig. 2). In Arabidopsis, root hairs form in alternating patterns, so that every root hair forming cell (trichoblast) is adjacent to a non-hair cell (atrichoblast). We quantified ROS accumulation by analyzing PO1 signal after confocal imaging, by drawing a line across the width of the root that spans 5 epidermal cell files so that the PO1 signal in 3 trichoblasts (numbered 1, 3, and 5) and 2 atrichoblasts (numbered 2 and 4) was quantified (Fig. 2A).

**Figure 2.**
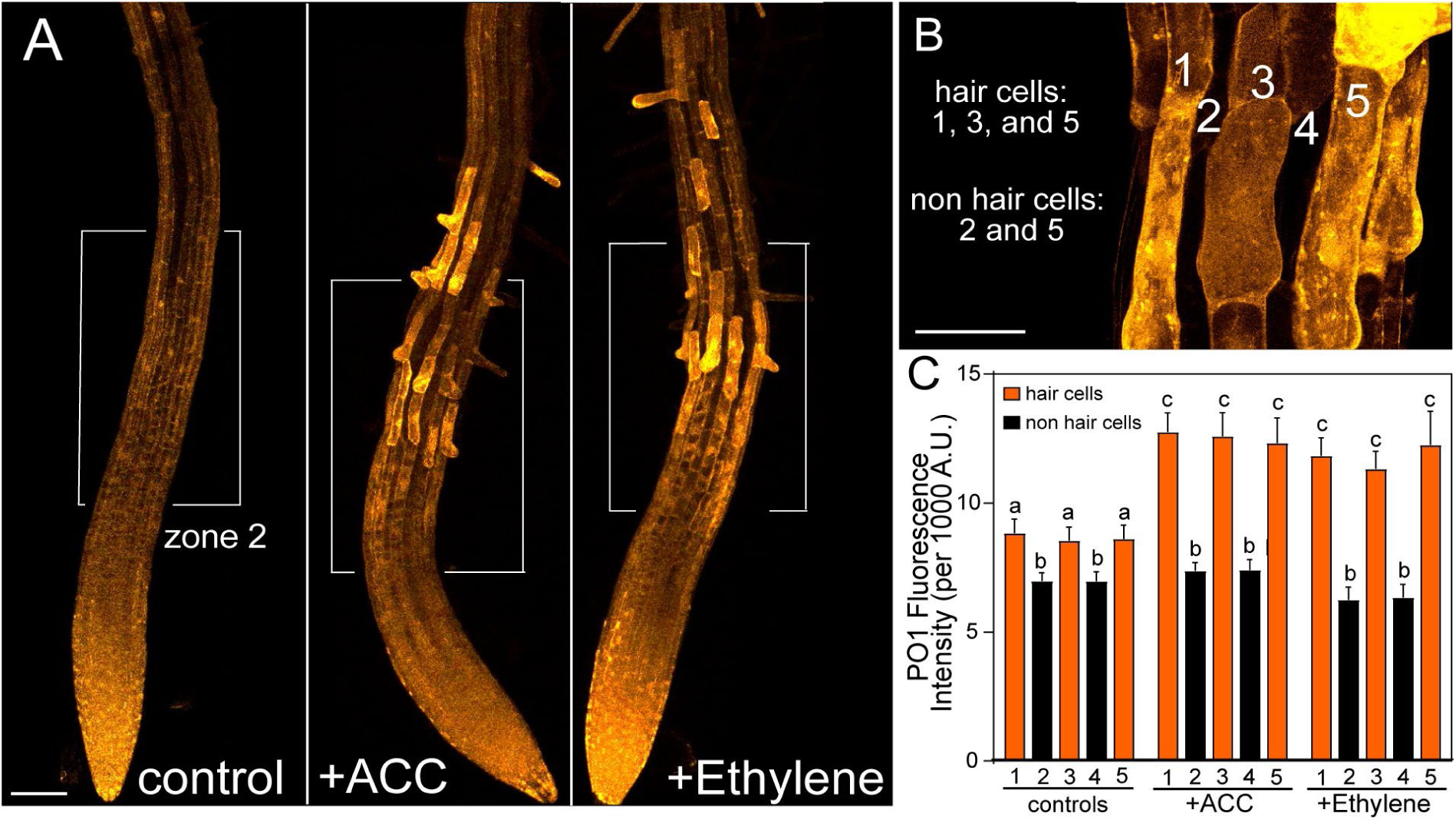
ACC and ethylene gas increase H_2_O_2_ accumulation in trichoblasts after 4 hours. (A) Representative roots illustrating the alternating PO1 fluorescence between trichoblast and atrichoblast epidermal cell files of cells in Col-0 roots with and without ACC and ethylene gas treatment for 4 hours. White brackets indicate zone 2 of the root. Scale bar is 100 µm. (B) Inset of hair cells (1, 3, 5) vs. non hair cells (2, 4) from roots treated with ethylene. Scale bar is 50 µm. (C) Quantification of PO1 fluorescence intensity in trichoblasts (orange bars: 1, 3 and 5) and atrichoblasts (black bars: 2, 4). Data are the means ± SEM of 3 independent experiments (n=18-24 seedlings/experiment). Columns with different letters indicate statistical significance compared to Col-0 untreated cells as determined by two-way ANOVA followed by Tukey’s multiple comparisons test.

We found that PO1 accumulation in trichoblasts in zone 2 of untreated seedlings was slightly, but not significantly higher than atrichoblasts. In trichoblast cells of ACC and ethylene treated seedlings, there was a significant increase in PO1 accumulation compared to trichoblasts of untreated seedlings. In contrast, there were no changes in PO1 in the atrichoblasts suggesting that ethylene and ACC treatment increased ROS levels in only the cells that formed root hairs, with PO1 fluorescence intensity values in trichoblasts after both treatments showing the same magnitude increase as roots treated with ACC (Fig. 2B). These results are consistent with this short term and low dose treatment with ACC leading to efficient conversion to ethylene that in turn produces elevated ROS in trichoblast cells (Fig. 2B). This is in contrast with some studies in which other developmental processes are altered by ACC acting directly, rather than conversion to ethylene (Li et al., 2021).

### ROS accumulation increases prior to root hair emergence

To determine whether ethylene-induced ROS accumulation drives root hair emergence, we asked whether ACC-induced ROS increases were detectable prior to the first ACC-induced root hair initiation. Wild type Col-0 seedlings were treated with ACC for either 2 or 4 hours and PO1 fluorescence was visualized in trichoblasts that did not have a root hair bulge (stages below +1) (Fig. 3A). This PO1 signal is only reported in the 2-hour ACC treatment, as there were insufficient number of cells without root hair bulges to quantify in root treated for 4 hours. Total PO1 accumulation was measured in 5 individual trichoblasts per root treated with and without ACC. Signal was quantified across the area of the entire cell and the average PO1 intensity of 24-30 individual cells was reported (Fig. 3B). Atrichoblast signal was not quantified as ROS levels did not change in those cells (Fig. 2B). These data showed a 1.3-fold increase in PO1 accumulation in hair cells of seedlings treated with ACC for 2h as compared to untreated controls, when trichoblast cells that had not yet begun to initiate root hairs were examined. These data are consistent with the hypothesis that ROS acts as a driver of root hair initiation downstream of ethylene signaling.

**Figure 3.**
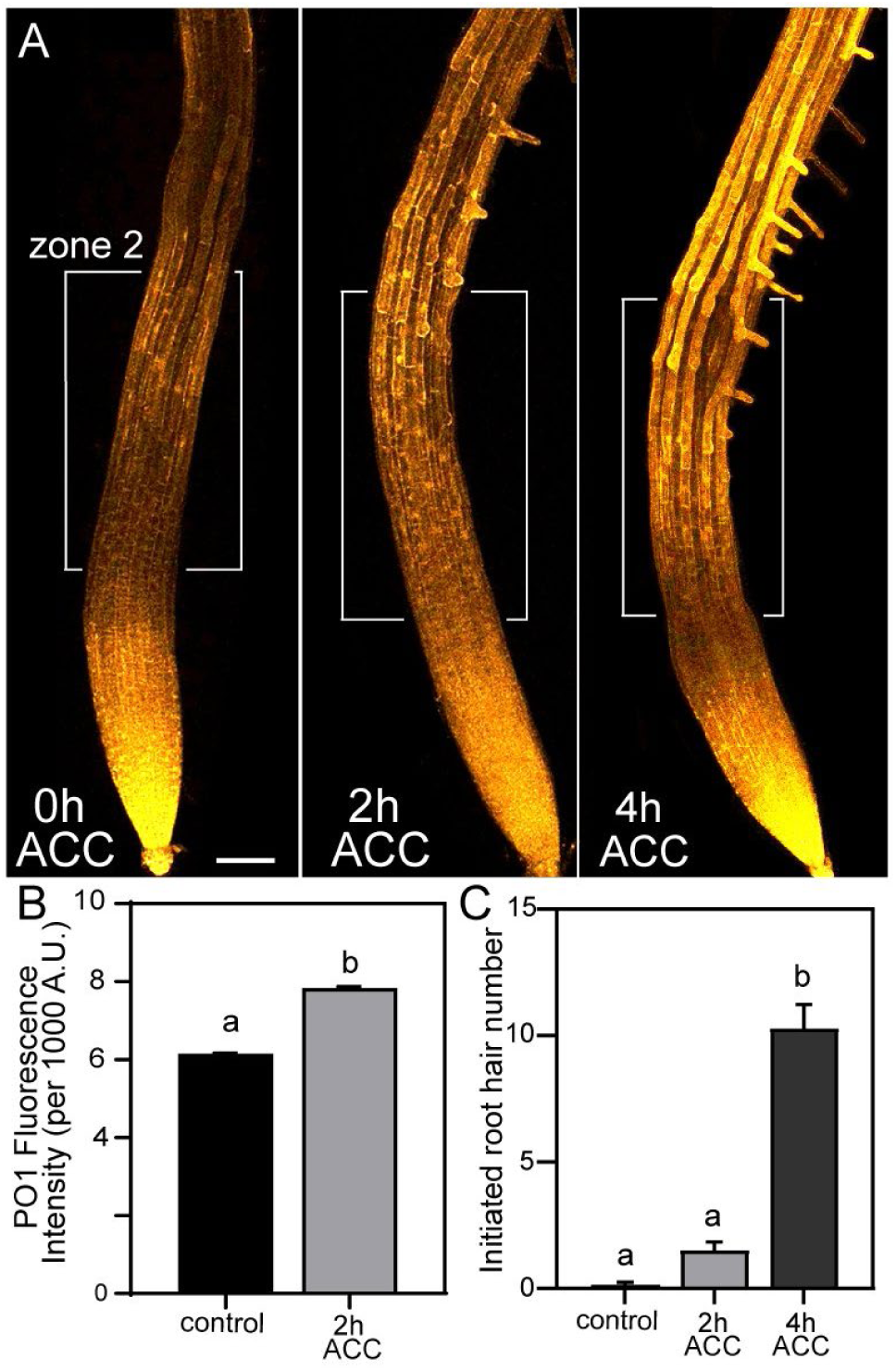
ACC-increased ROS accumulation precedes root hair initiation. (A) Representative images of epidermal PO1 fluorescence in untreated Col-0 and Col-0 treated with ACC for 2 or 4h. Scale bar is 100 µm. White brackets indicate zone 2 of the root. (B) Quantification of trichoblasts that had not yet formed root hair bulges of control and 2h ACC treated seedlings. (C) Quantification of number of initiated root hairs in zone 2 of the root (500 µm-1000 µm from root tip) in untreated and 2 and 4h ACC-treated seedlings. Data are mean ± SEM of individual cells from 3 independent experiments (n= 12-18 seedlings/experiment). Columns with different letters indicate statistical significance compared to untreated hair cells as determined by Student’s t-test (p < 0.0001) in B. Columns with different letters indicate statistically significant differences determined by one-way ANOVA followed by Tukey’s multiple comparisons test in C.

### Ethylene signaling mutants show altered ROS accumulation patterns and root hair phenotypes

The ethylene signaling pathway is well defined (Fig. 4A) and mutants in key signaling proteins, including receptors and transcription factors are available. ROS accumulation was examined in response to ACC treatment in these ethylene signaling mutants to ask whether this response was dependent on the ethylene signaling pathway and downstream transcriptional responses. The number of root hairs and average length were previously reported in loss-of-function and gain-of-function ethylene receptor mutants, *etr1-7 and etr1-3*, and the transcription factor mutant *ein3eil1* in the presence and absence of ACC treatment (Harkey et al., 2018). *etr1-7* is a LOF mutant in which the ETR1 receptor is inactive, therefore the ethylene signaling pathway is constitutively signaling and there are increased numbers of root hairs independent of ACC addition, while *etr1-3* is a GOF mutant in which the ETR1 receptor is always active leading to the ethylene signaling pathway being inhibited (Harkey et al., 2018; Hua and Meyerowitz, 1998). The double mutant *ein3eil1* has mutations in genes encoding EIN3 and EIL1 TFs (Chao et al., 1997; Solano et al., 1998). Both *etr1-3* and *ein3eil1* have reduced root hair initiation in response to ACC treatment (Harkey et al., 2018). We examined the PO1 distribution patterns in root hair cells with and without ACC treatment in these three mutants.

**Figure 4.**
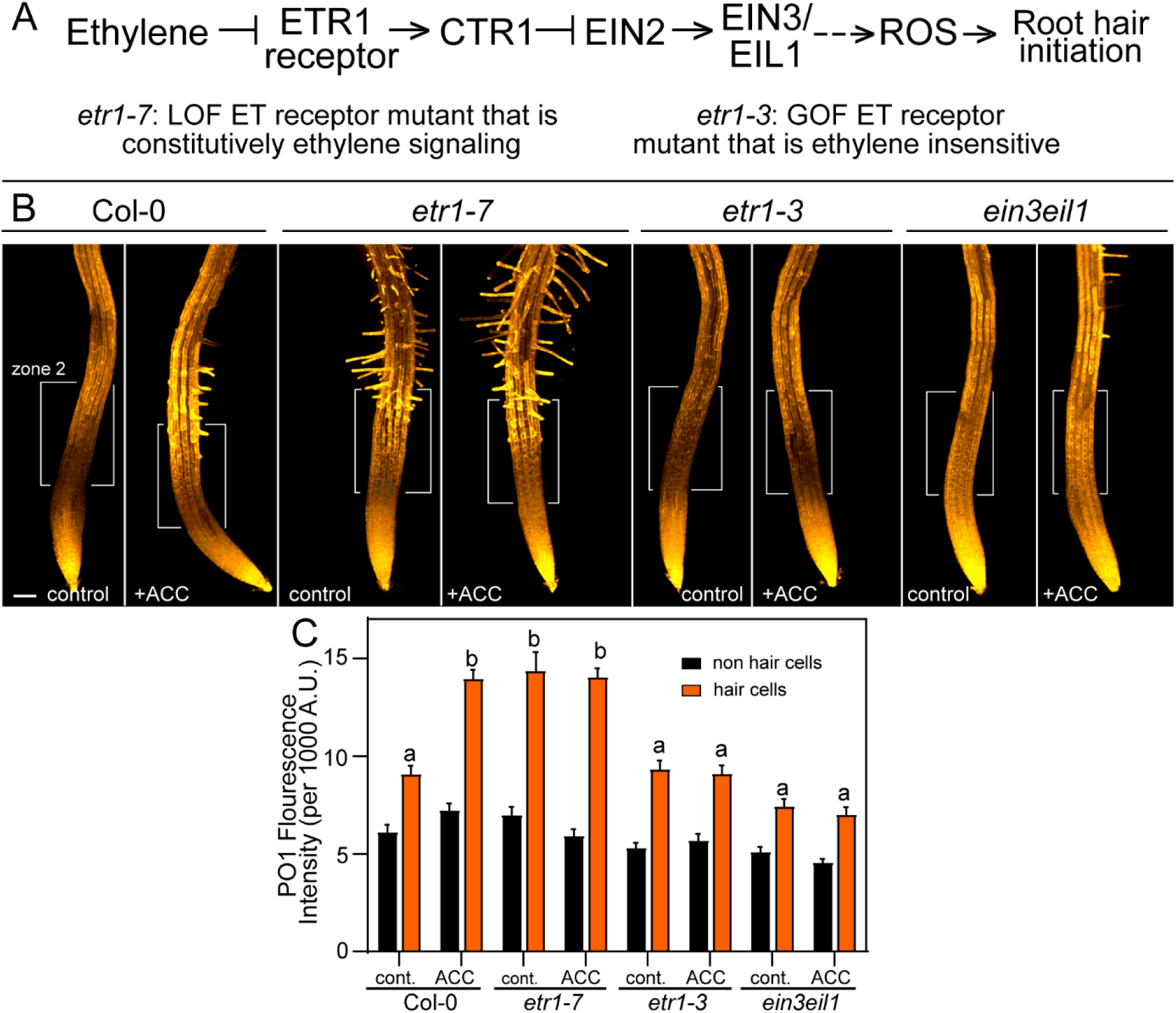
The ETR1 receptor and EIN3/EIL1 transcription factors are required for ethylene-induced ROS accumulation and root hair proliferation. (A) A schematic diagram of the ethylene signaling pathway and explanation of the character of the *etr* mutants. ETR1 (ethylene resistant 1) is the ethylene receptor controlling root hair formation, CTR1 (constitutive triple response 1 is a kinase, EIN2 (ethylene insensitive 2) is a signaling protein, EIN3 and EIL1 (EIN3-like 1) are transcription factors. (B) Representative images of PO1 epidermal fluorescence in Col-0, *etr1-7, etr1-3*, and *ein3-1eil1-1* treated with and without ACC for 4 hours. 18-20 seedlings from each genotype and treatment were imaged. Scale bar is 100 µm. (C) Quantification of PO1 fluorescence intensity in hair cells and non-hair cells. For each root, PO1 fluorescence was quantified in two hair cells and two non hair cells and the values were pooled to obtain this average. Data are means ± SEM of 3 experiments (n=18-20 seedlings/experiment). Columns with different letters indicate statistical significance in PO1 signal compared to other hair cells as determined by two-way ANOVA followed by Tukey’s multiple comparisons test.

We visualized PO1 fluorescence via LSCM and saw that the constitutive ethylene signaling mutant *etr1-7* had increased H_2_O_2_ in root hair cells and an increased number of trichoblasts with emerged root hairs regardless of ACC treatment (Fig. 4B). The PO1 signal in *etr1-7* in the absence of ACC is significantly elevated over untreated Col-0, but is equivalent to ACC treated Col, and is not significantly changed by ACC treatment. The opposite response was seen in the ethylene-insensitive *etr1-3* and *ein3eil1* mutants, as they showed no change in PO1 signal in response to ACC treatment, resulting in a significantly lower level than ACC treated Col-0 and no induction of root hair initiation. Together, these results suggest that the ETR1 receptor and EIN3/EIL1 TFs are required for ACC-induced ROS accumulation and root hair initiation and suggesting that in these experiments ACC is acting via conversion to ethylene.

### Mutants with altered root hair formation also show altered ROS accumulation

We also examined the effect of mutations that alter root hair formation on ROS levels. The *werewolf* (*wer*) mutant has increased root hair formation, while the *caprice* (*cpc*) mutant with decreased root hair formation. ROS accumulation was significantly higher in root hairs along the root in *wer* compared to both Col-0 and *cpc*, while *cpc* had significantly less ROS accumulation along the root compared to Col-0 (Fig. 5).

**Figure 5.**
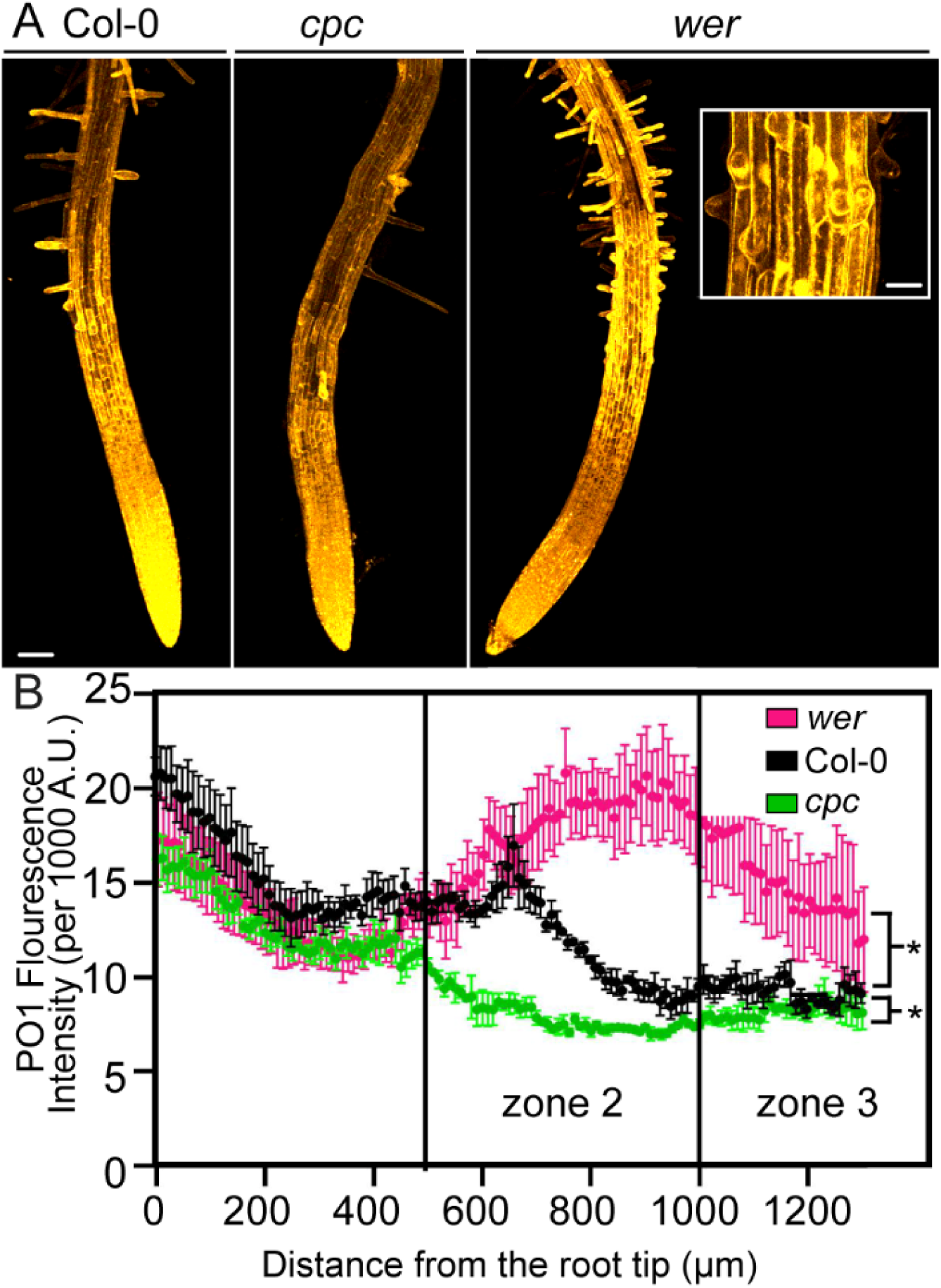
ROS accumulation increases in *werewolf* and decreases in *caprice* compared to Col-0. (A) Representative images of 5-day old untreated Col-0, *cpc*, and *wer* roots stained with PO1. Scale bar is 100 μm. Inset is 40x image of *wer* root to illustrate ectopic root hair formation. Scale bar = 25 μm (B) Quantification of total PO1 accumulation along the root starting approximately 200 μm from the root tip and ending at 1300 μm. This region encompasses both zones 2 and 3. Data are means ± SEM and representative of 2 independent experiments (9-15 seedlings per experiment). Asterisks indicate statistically significant differences as determined by One-way ANOVA.

### The RBOHC knockout mutant, *rhd2-6*, shows decreased ROS accumulation after ACC treatment

It has been previously reported that the respiratory burst oxidase homolog C (RBOHC) is involved in ROS production and subsequent root hair elongation (Foreman et al., 2003). We have also shown that there is decreased ROS in root hairs of the *rhd2-6* mutant, which has an insertion mutation in the *RBOHC* gene (Gayomba and Muday, 2020). Therefore, we asked whether RBOHC is required for ACC-induced ROS increases and root hair initiation. We examined root hair numbers via light microscopy in ACC-treated *rhd2-6* (*rbohc*) seedlings (Fig. 6). Root hair number in the ACC-treated *rhd2-6* mutant was significantly less than ACC-treated Col-0, suggesting that RBOHC contributes to ethylene-induced root hair initiation (Fig. 6B). We also quantified the length of all root hairs in Zone 2 using the segmented line tool in ImageJ. A histogram showing the distribution of lengths of root hairs (reported as percentage in each length bin out of total number of roots) revealed that the length of root hairs increased after ACC treatment and that the *rhd2-6* mutant has reduced root hair elongation in the presence of ACC relative to Col-0.

**Figure 6.**
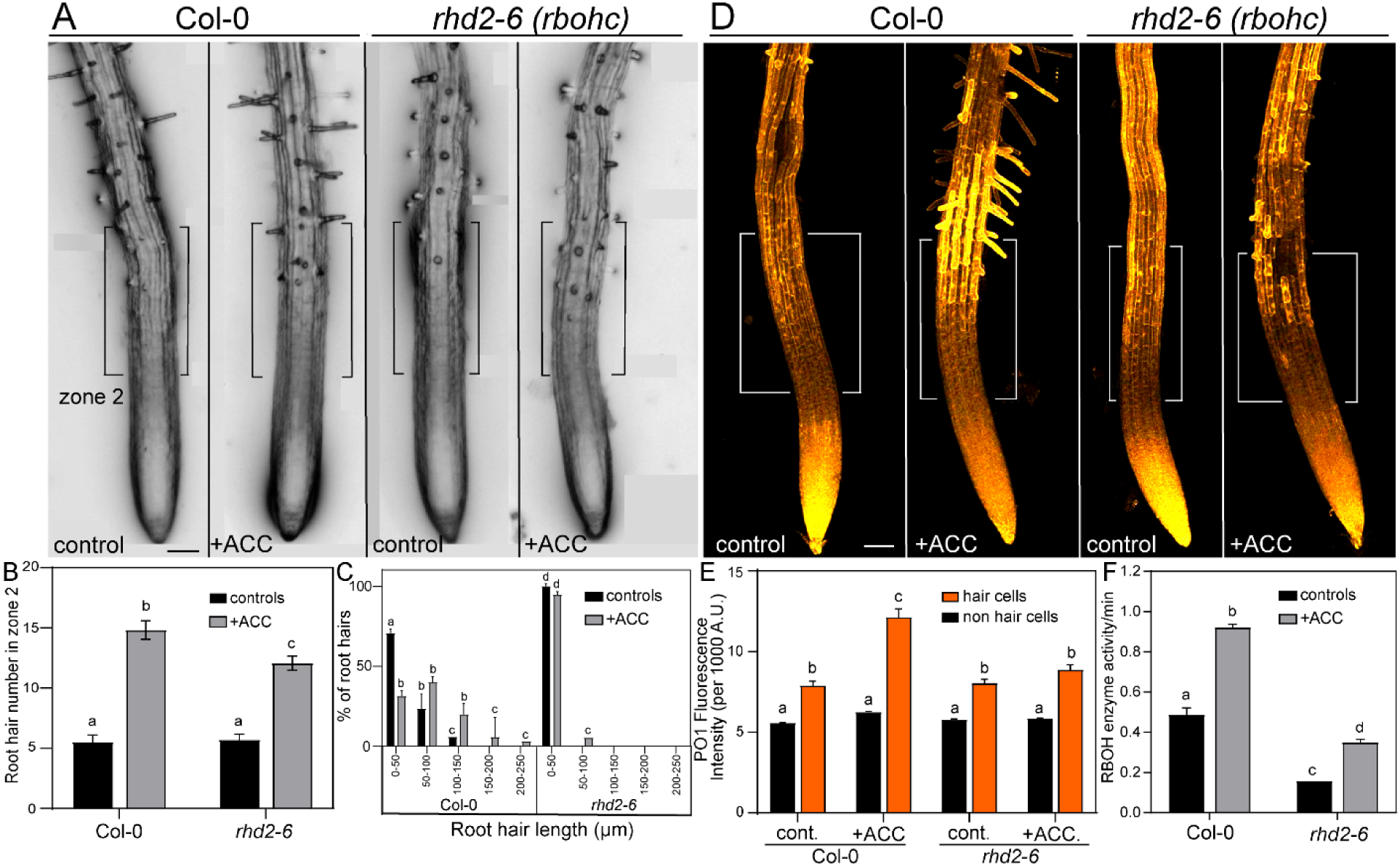
RBOHC activity contributes to ACC-induced ROS accumulation. (A) Representative images of root hairs of Col-0 and *rhd2-6* treated with and without ACC for 4 hours. Black brackets indicate zone 2 of the root. (B) Root hair quantification of untreated and 4h ACC treated seedlings. Data are means ± SEM of total RH in zone 2 from 3 experiments (n=18-24 seedlings per experiment). Columns with different letters indicate statistically significant differences determined by two-way ANOVA followed by Tukey’s multiple comparisons test. (C) Quantification of percent of root hairs within a certain micron length of Col-0 and *rhd2-6* treated with and without ACC. Columns with different letters indicate statistically significant differences determined by two-way ANOVA followed by Tukey’s multiple comparisons test. (D) Representative images of PO1 epidermal fluorescence in Col-0 and *rhd2-6* treated with and without ACC for 4 hours. White brackets indicate zone 2 of the root. (E) Quantification of PO1 fluorescence intensity in hair cells and non-hair cells. For each root PO1 fluorescence was quantified in two hair cells (cells 1 and 5 from Figure 2) and two nonhair cells (cells 2 and 4 from Figure 2) and the values pooled to obtain this average. Data are the means ± SEM of 3 experiments (n=18-24 seedlings/experiment). Columns with different letters indicate statistical significance in PO1 signal compared to other hair cells as determined by two-way ANOVA followed by Tukey’s multiple comparisons test. (F) Quantification of RBOH activity as reported as changes in formazan concentration per minute in roots of Col-0 and *rhd2-6* treated with and without ACC for 4 hours. Data are means ± SEM of 3 experiments. Columns with different letters indicate statistical significance as determined by two-way ANOVA followed by Tukey’s multiple comparisons test. Scale bars for all images are 100 µm.

We also visualized the PO1 fluorescence by LSCM in the *rhd2-6* mutant with and without ACC treatment (Fig. 6D). We report the average of 2 trichoblast cells (1 and 5 from Fig. 2) and two atrichoblast cells (2 and 4) in Fig. 6E. This reveals a significant increase in H_2_O_2_ fluorescence accumulation in Col-0 hair cells, while in *rhd2-6* there is no significant increase after ACC treatment (Fig. 6E), suggesting that RBOHC contributes to ethylene-induced ROS accumulation. The data is reported for all 5 cell files in Fig.s S2. These images reveal slight increases in numbers of root hairs in ACC-treated *rhd2-6* roots, suggesting that the ACC also acts to induce root hairs in an RBOHC-independent mechanism.

As there are also other RBOH enzymes expressed in roots, we asked whether these other enzymes affect root hair formation. RBOHD and RBOHF are highly expressed in roots and have been shown to be involved in regulation of root elongation and lateral root development (Chapman et al 2019). Therefore, to determine these other root expressed RBOHs contributed to ethylene-induced ROS accumulation, PO1 accumulation patterns were examined in the *rbohd/f* double mutant and compared to *rhd2-6*. No change was seen in ROS levels in hair cells of *rbohd/f*, suggesting that these RBOHs do not contribute to ethylene-induced ROS synthesis in root hair cells (Fig. S2).

### RBOH transcript abundance and enzyme activity increases in response to ethylene signaling

Elevated ethylene may either increase *RBOHC* transcript abundance or enzyme activity. We examined several transcriptomic datasets with ACC or ethylene treated roots and found a subtle change in the *RBOHC* transcripts, but not significant enough to pass the filtering on this transcriptomic analysis (Harkey et al., 2018; Harkey et al., 2019) (Table S1). However, we also performed qRT-PCR in Col-0 and *ein3eil1* treated with and without ACC to examine changes in abundance in the transcript encoding RBOHC. *RBOHC* transcript abundance was 2-fold higher in Col-0 treated with ACC compared to Col-0 controls. The levels of *RBOHC* transcript abundance in untreated *ein3eil1* were equivalent to Col-0, while and the increase by ACC treatment in Col-0 was lost in *ein3eil1* (Fig. S3B). To determine if ethylene signaling regulated RBOH enzyme activity, a spectrophotometric assay using nitro blue tetrazolium (NBT) dye as an electron acceptor, was performed. NBT is reduced by superoxide to monoformazan and this reduction can be detected at 530 nm. This experiment was performed in protein extracts from 7-day old roots of Col-0 and *rhd2-6* treated with and without ACC for 4 hours. Older roots were used in this experiment to obtain an adequate amount of protein for these assays. There was a significant 2-fold increase in monoformazan production in protein extracts of Col-0 roots treated with ACC for 4 hours compared to controls (Fig. 6F). The enzyme activity of roots of *rbohc/rhd2-6* was 3-fold lower in both control and ACC treated conditions than roots of similarly treated Col-0 (Fig. 6F). These data are consistent with previous results indicating that the RBOHC enzyme constitutes the majority of the RBOH activity in roots (Chapman et al., 2019; Gayomba and Muday, 2020).

### Transcription of *PRX44* increases with ACC treatment driving ROS accumulation and ethylene-induced root hair initiation

The *ein3/eil1* transcription factor mutant has reduced root hair initiation and a reduction in PO1 signal in trichoblasts suggesting that there is transcriptional regulation of ROS producing enzymes that drive root hair initiation. We examined a previously published microarray time course experiment in roots treated with ACC (Harkey et al., 2018), to identify candidate transcriptional targets profiling the expression pattern of transcripts encoding both ROS producing enzymes and proteins linked to trichoblast cell specification. ROP2, ROPGEF3, ROPGEF4, GL2, RSL4 and RHD6, which are linked to root hair initiation (Denninger et al., 2019), showed no transcriptional response to ACC treatment and primers that recognize RSL1 and RSL2 are not present on the microarray and could not be examined in this dataset (Table S1). This transcriptomic analysis did not identify the previously reported ethylene induction of RSL4 (Feng et al., 2017), which was revealed with high dose treatment of ethylene that led to ectopic root hair formation from non-hair cells, suggesting the RSL4 transcript changes may be dose dependent or ethylene specific. A number of transcripts encoding class III peroxidases (PRXs) changed in abundance in response to ACC, including the transcript encoding *PRX44*, which increased by 3-fold. Class III peroxidases are specific to plants and exist in large multigene families. They have been implicated in root hair tip growth, as null mutants have shorter root hairs compared to wild type (Mangano et al., 2017). Recent work has shown that auxin induces expression of genes encoding four class III peroxidases (PRX), which results in an increase in both root hair length and ROS accumulation (Mangano et al., 2017), however, the role of ethylene signaling in regulating class III PRX expression to modulate root hair initiation or elongation has not been reported.

We examined control and ACC-treated seedlings harboring the *PRX44* promoter driving GFP and examined GFP expression via LSCM. Seedlings treated with ACC showed a statistically significant >2-fold increase in GFP fluorescence, suggesting that the *PRX44* promoter is induced downstream of ethylene signaling (Fig. 7A-B). qRT-PCR was also performed in Col-0 and *ein3eil1* treated with and without ACC to examine changes in abundance in the transcript encoding PRX44. We found that there was a 2-fold increase in *PRX44* transcript abundance in Col-0 treated with ACC compared to Col-0 controls. We also found that there was a significant decrease in *PRX44* transcript abundance in untreated and ACC treated *ein3eil1* compared to Col-0 (Fig. S3A). We also asked whether these peroxidases participate in ACC-regulated root hair initiation. We examined root hair numbers via light microscopy in Col-0 and *prx44-2* seedlings treated with and without ACC. The number of root hairs formed in the *prx44-2* mutant treated with ACC was significantly less than in ACC treated Col-0 (Fig. 7C-D). The root hairs in the ACC-treated mutant were also significantly shorter compared to Col-0, which is consistent with the phenotype that has been reported in response to auxin treatment (Fig. 7E) (Mangano et al., 2017). When PO1 accumulation patterns were examined via LSCM, we observed no significant increase in PO1 accumulation in hair cells of *prx44-2* seedlings treated with ACC (Fig. 7F-G). We also examined *prx73-4*, since transcripts encoding PRX73 have been reported to be induced with auxin treatment to drive root hair elongation (Mangano et al., 2017) and these transcripts increased in abundance in response to ACC treatment (Table S1). When treated with ACC, the *prx73-4* mutant exhibited root hair initiation and ROS accumulation patterns that were similar to wild-type seedlings, suggesting that this class III PRX is not involved in ethylene-induced ROS synthesis and root hair initiation. These combined data indicate that both the class III PRX44 and PRX73 are transcriptionally regulated by ethylene, but only PRX44 contributes to ethylene induced ROS accumulation in root hairs and root hair initiation.

**Figure 7.**
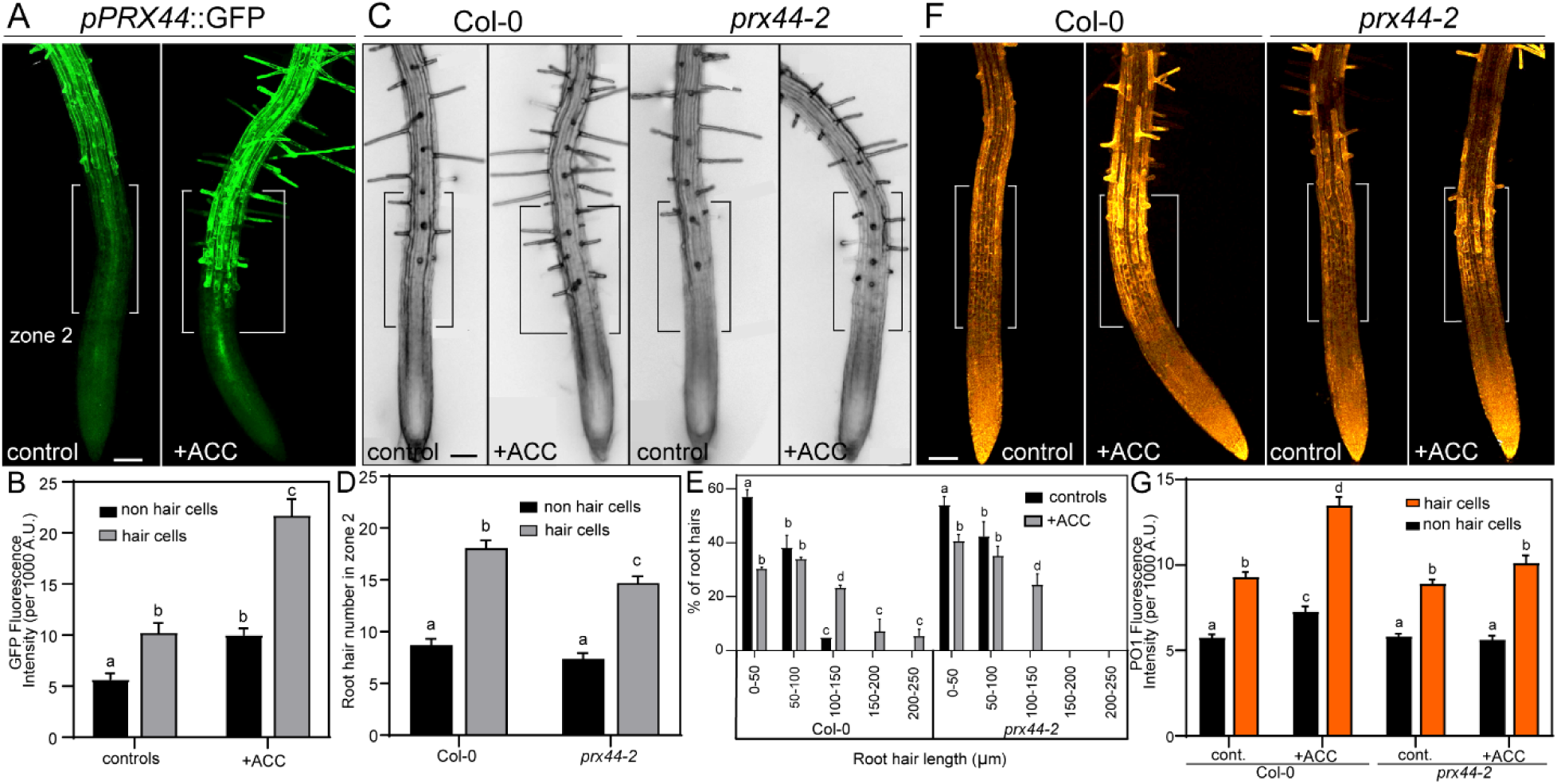
*pPRX44*::GFP expression increases in roots treated with ACC and PRX44 contributes to ethylene-induced root hair formation and H_2_O_2_ accumulation in root hair cells. (A) Representative images of *pPRX44*::GFP treated with and without ACC for 4 hours. White brackets indicate zone 2 of the root. (B) Quantification of GFP signal in hair and non-hair cells of *pPRX44*::GFP treated with and without ACC. Data are means ± SEM (C) Representative images of RH number in Col-0 and *prx44-2* treated with and without ACC for 4 hours. Black brackets indicate zone 2 of the root. (D) Root hair quantification of untreated and 4h ACC treated seedlings. Data are means ± SEM. of 3 independent experiments (n=18-24 seedlings per experiment). Columns with different letters indicated statistically significant differences determined by two-way ANOVA followed by Tukey’s multiple comparison test. (E) Quantification of percent of root hairs within a certain length for Col-0 and *prx44-2* treated with and without ACC. Columns with different letters indicate statistically significant differences determined by two-way ANOVA followed by Tukey’s multiple comparisons test. (F) Representative images of Col-0 and *prx44-2* treated with and without ACC for 4 hours and stained with PO1. White brackets indicate zone 2 of the root. (G) Quantification of PO1 accumulation in hair and non-hair cells of Col-0 and *prx44-2* treated with and without ACC for 4 hours (n=12-18 seedlings per experiment). For each root PO1 fluorescence was quantified in two hair cells (cells 1 and 5 from Figure 2) and two nonhair cells (cells 2 and 4 from Figure 2) and the values pooled to obtain this average. Data are means ± SEM of 3 experiments. Columns with different letters indicate statistical significance compared to all other hair cells as determined by two-way ANOVA followed by Tukey’s multiple comparisons test. Scale bars for all images are 100 µm.

## DISCUSSION

Ethylene is a key hormonal regulator of root hair initiation and elongation (Guzman and Ecker, 1990; Kieber et al., 1993). Although the proteins that drive ethylene signaling are well characterized, the downstream proteins that control the root hair developmental processes have received less attention. Reactive oxygen species (ROS) are critical for both root hair initiation and elongation (Foreman et al., 2003; Gayomba and Muday, 2020). In root hairs, one source of ROS is the NADPH oxidase (NOX)/respiratory burst oxidase homolog C (RBOHC) that is localized to root hairs to facilitate cell wall loosening and subsequent tip focused Ca^2+^ accumulation leading to root hair cell elongation (Foreman et al., 2003; Monshausen et al., 2007). The plant hormone auxin also increases ROS accumulation to drive root hair elongation (Mangano et al., 2017), suggesting that ROS could also act as a signaling molecule in ethylene-induced root hair initiation and elongation (Foreman et al., 2003; Gayomba and Muday, 2020; Jones et al., 2007; Takeda et al., 2008). In this study, we examined the effects of ethylene signaling on root hair initiation and ROS accumulation to determine if increased ethylene drives root hair initiation through regulation of ROS producing enzymes and subsequent ROS accumulation in trichoblasts from which root hairs form.

We examined root hair number and ROS accumulation in Col-0 treated with low concentrations of ethylene gas (0.5 ppm) or the ethylene precursor, ACC (0.7 µM), for 4 hours. There was an increase in the number of trichoblasts that formed root hairs in ethylene and ACC treated seedlings compared to untreated controls, which was most pronounced in a 500 µm region starting 500 µm from the root tip, which we call zone 2. Using a ratiometric reporter of ROS-induced gene expression, ZAT12p-ROS, we found that in ACC-treated roots, ROS dependent gene expression was increased in zone 2 in ACC-treated roots compared to untreated controls. We also examined the effect of ethylene and ACC on hydrogen peroxide accumulation using a boronate based hydrogen peroxide selective sensor, PO1. Both ethylene and ACC lead to elevated H_2_O_2_ accumulation in the trichoblast cells in zone 2 after 4 hours of treatment, but PO1 signal was at constant lower level in atrichoblast cells. There was a significant increase in PO1 signal within 2 hours after ACC treatment in cells that had not yet formed root hair bulges, suggesting that ethylene-induced ROS acts to drive root hair initiation.

To examine whether canonical ethylene signaling controls ACC-dependent ROS synthesis driving root hair formation, we examined these responses in several ethylene signaling mutants. In the ethylene insensitive *ein3eil1* and *etr1-3* mutants the effect of ACC treatment on ROS synthesis and root hair formation is lost. In contrast, the *etr1-7* mutant allele that has constitutive signaling, has increased ROS and root hair formation. These findings are consistent with our prior reports that showed that effects of this ACC dose and under these growth conditions on root elongation, gravitropism, and lateral root formation are lost in the ethylene insensitive *ein2-5* and *etr1-3* mutants (Harkey et al., 2018; Lewis et al., 2011; Negi et al., 2010). These findings contrast with other processes in which higher doses of ACC can affect development without conversion to ethylene, especially when the ACC oxidase enzyme is limited (Li et al., 2021; Polko and Kieber, 2019; Van de Poel, 2020; Vanderstraeten et al., 2019). We also do not see ectopic root hair formation in response to these short term and low dose treatments with ACC and ethylene, as reported recently when seedlings were treated with 10ppm ethylene for 2 days (Qiu et al., 2021), which is a 12-fold longer and 20-fold higher concentration.

If ROS is driving root hair formation, then it is predicted that mutants with impaired root hair initiation, may have altered ROS levels. We examined the PO1 signal in the root hairless mutant, *caprice (cpc)*, and the *werewolf (wer)* mutant that forms ectopic root hairs, both of which have mutations in MYB-like DNA binding proteins (Lee and Schiefelbein, 1999; Wada et al., 1997). The *cpc* mutant showed a significant decrease in PO1 signal when it is measured using line profiles along the root while *wer* showed a significant increase, suggesting that mutants with altered root hair formation also show altered ROS accumulation patterns. Similarly, the *rhd2-6* mutant, which was isolated for altered root hair elongation (Foreman et al., 2003) and has impaired root hair formation (Gayomba and Muday, 2020) also had reduced levels of ROS, suggesting a positive relationship between root hair formation and ROS levels using root hair mutants.

Based on previous work detailing the role of RBOHC in root hair initiation and growth (Foreman et al., 2003; Gayomba and Muday, 2020), we also asked whether this enzyme was contributing to ethylene-induced ROS accumulation and subsequent root hair initiation and/or elongation. ACC treated *rhd2-6*, which is an *RBOHC* null mutant, had a reduced number of root hairs, less root hair elongation, and decreased ROS accumulation in trichoblasts compared to Col-0. We also saw more than a 2-fold increase in RBOH activity in response to ACC treatment and a 3-fold reduction in activity in *rhd2-6* (*rbohc*). These data suggest that ethylene signaling induces ROS production in roots via increases in RBOHC enzyme activity to drive root hair initiation and elongation but could also come from enhanced RBOHC synthesis. We examined several transcriptomic datasets (Harkey et al., 2018; Harkey et al., 2019) and found evidence of changes in *RBOHC* transcripts in response to ACC treatment, which were confirmed by qRT-PCR. The ACC-induction was lost in the ein3/eil1 mutant.

To determine if other root hair specific proteins and/or ROS producers are regulated by ethylene, we widened our analysis of these previously published ACC and ethylene transcriptomic datasets. Although ACC did not substantially change the abundance of transcripts encoding RHD2 (RBOHC), ROP2, ROPGEF3, ROPGEF4, GL2, RHD6, or RSL4, we identified several transcripts that showed substantial increase in response to ACC. Of particular interest were increases in transcripts encoding two class III peroxidase enzymes PRX44 and PRX73, which showed a significant increase in abundance after 4 hours of ACC treatment. ACC treatment of seedlings harboring a *pPRX44*::GFP construct lead to significant increases in GFP fluorescence intensity in zone 2 compared to untreated controls, consistent with transcriptional regulation of this gene. We also performed qRT-PCR and found a 2-fold increase in PRX44 transcripts in response to ACC treatment. Additionally, this ACC effect was lost in the *ein3/eil1* mutant and that the abundance of *PRX44* transcripts was significantly lower in *ein3/eil1* even the absence of ACC treatment.

ACC treatment of the null mutants *prx44-2* led to decreased ROS accumulation in root hair forming trichoblasts, while there was no effect in *prx73-4* as compared to Col-0. The decreased PO1 signal in *prx44-2* was accompanied by significant reductions in the number of root hairs formed in this same root region and the length of these root hairs. Together these experiments implicate RBOHC and Class III peroxidase enzymes in ethylene-regulated ROS accumulation and root hair initiation.

These class III peroxidase enzymes have also been implicated as regulators of root hair growth in response to environmental cues. Two PRXs are involved in auxin-dependent root hair elongation, as null mutants treated with auxin have shorter root hairs and less ROS compared to wild type (Mangano et al., 2017). Another report demonstrated that PRX enzymes function to modify the localization of extension proteins to control cell wall structure to allow increased elongation (Marzol et al 2020). Recent work has also indicated a role for PRXs as positive regulators of root hair growth at low temperatures (Pachecho et al 2022). These results showed that low temperature induced root hair growth required peroxidase enzyme activity and upregulated expression of PRXs that modulate ROS homeostasis and extensin stabilization to drive root hair growth (Pacheco et al 2022). Prior publications and these results are consistent with a role for peroxidases as drivers of root hair development and modulation of the apoplastic ROS pool.

To summarize our findings on ethylene-dependent root hair initiation via increased ROS synthesis and to tie these findings with other studies, we present a model in Fig. 8. Low dose and short 4-hour treatments with ACC led to efficient conversion of ACC to ethylene by roots to initiate an ethylene signaling cascade. Ethylene acts through the ETR1 receptor and the canonical signaling pathway to increase the activity of EIN3/EIL1 TFs. EIN3/EIL1 binds to the root hair specific TF RHD6 to induce RSL4 expression and root hair elongation (Feng et al., 2017). Root hair growth is controlled by several proteins leading to activation of the RHD6 TF. Downstream of RHD6, the RSL4 TF acts to define final root hair length based on its level of expression (Datta et al., 2015; Mangano et al., 2017). Previous work has shown that EIN3 physically interacts with RHD6 to form a transcriptional complex that coactivate RSL4 to promote root hair elongation (Feng et al., 2017), and that EIN3 and RSL4 both act to downregulate GL2 expression to drive ectopic root hair formation with long term treatments with high levels of ethylene (Qiu et al., 2021). It has also been shown that auxin treatment results in increased RSL4 transcript abundance, which binds to the promoters of RBOH and four class III peroxidase (*PRX*) genes, including *PRX44*, to drive ROS synthesis in root hair cells. RSL4 activation via auxin induces class III peroxidase and *RBOH* expression, which leads to ROS accumulation required to drive root hair elongation (Mangano et al., 2017). We find that ACC treatment increases activity of RBOHC enzymes leading to increased ROS and root hair elongation, without changing the abundance of *RBOHC* transcripts. Additionally, ACC treatment leads to the transcriptional regulation of class III peroxidases and RBOHC, resulting in ROS production and root hair initiation. Although the reduction in ROS in *rhd2-6* and *prx44*-2, in both the presence absence of ACC was expected, we did not expect that there would still be an increase in root hair number in response to ACC. This data suggests that perhaps the ROS selective dyes that we have access to are not sensitive enough to detect the small changes in ROS that occur prior to root hair initiation in these mutants or that there are additional mechanisms that may facilitate ethylene-induced root hair formation, including redundant action of these two enzymes. Further investigation regarding the mechanisms of ethylene-regulated ROS synthesis is required. For example, one important question to be answered is how ethylene signaling regulates RBOH activity. This could occur through a number of different mechanisms, such as calcium binding or phosphorylation, which are known to regulate RBOH enzymes (Postiglione and Muday, 2020).

**Figure 8.**
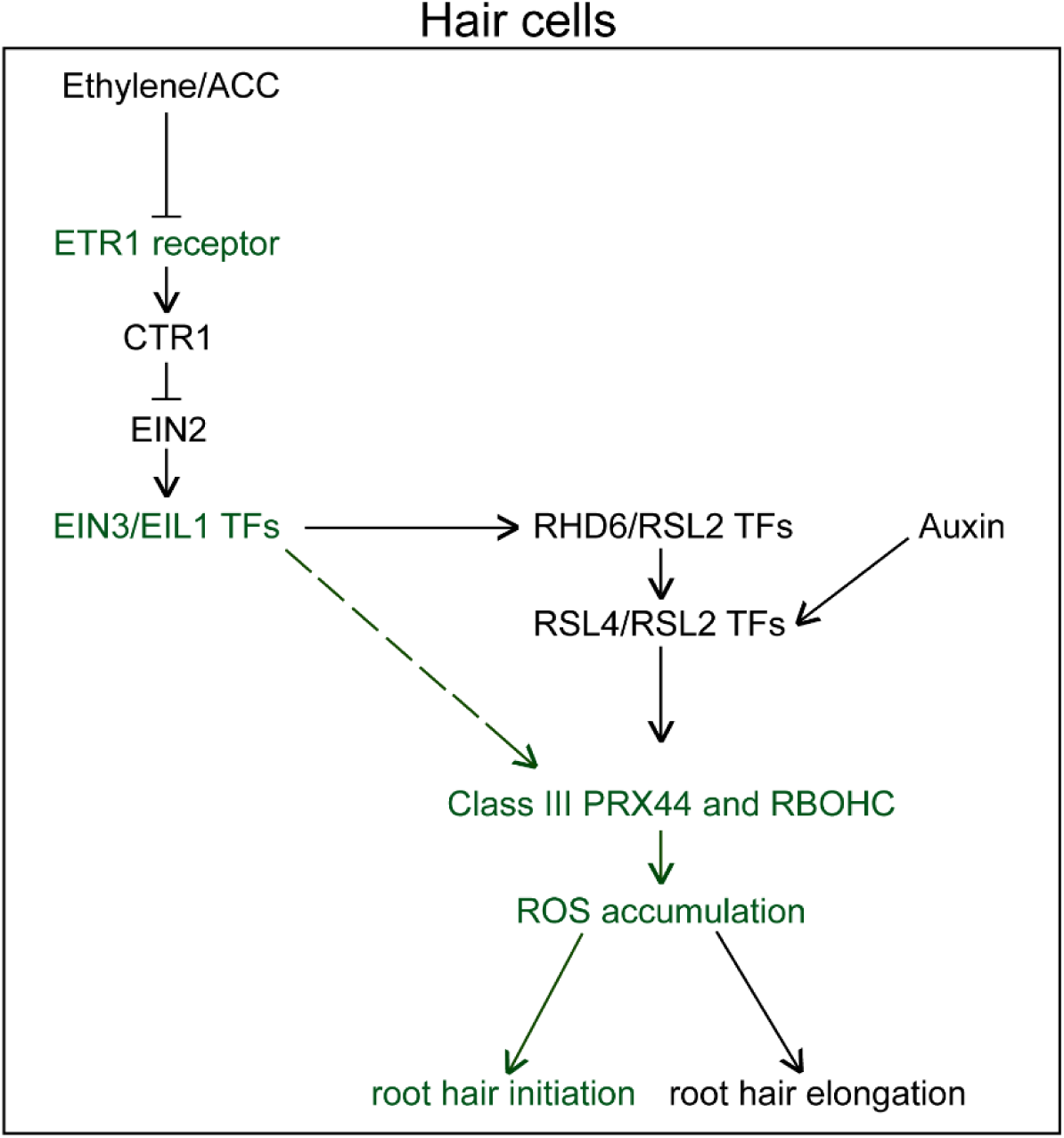
Summary of mechanisms by which ethylene and auxin modulate ROS and root hair formation. In hair cells, ethylene, acting through the ETR1 receptor and the canonical ethylene signaling pathway induces accumulation of EIN3/EIL1 TF proteins. Previous work has shown that EIN3/EIL1 physically interacts with RHD6/RSL2 TFs to induce expression of RSL4/RSL2 transcripts leading to root hair elongation (Feng et al., 2017). It has also been shown that RSL4/RSL2 is induced by auxin signaling and that RSL4/RSL2 binds to the promoters of RBOH and class III PRX genes to induce transcript expression and root hair elongation (Mangano et al., 2017). Here, highlighted in green, we have shown that ethylene signaling through ETR1 and EIN3/EIL1 induces ROS accumulation through RBOHC enzyme activity and PRX44 transcript expression to drive root hair initiation.

This work supports the role of ROS as a signaling molecule, in which ROS are produced in response to hormonal cues to drive developmental processes. We provide new evidence for the ethylene regulation of the ROS producing enzymes, RBOHC and class III PRX44. However, there are other ROS producing and scavenging enzymes, such as superoxide dismutase (SODs), that require additional study to understand the numerous developmental mechanisms that ROS may play a role in. It is also crucial to understand the molecular mechanisms by which ROS regulates the activity of proteins that drive root hair initiation and elongation. ROS can oxidize cysteine residues in proteins to change their conformation and activity (Couturier et al., 2013) so a critical next step is to identify those root hair proteins are modified by ROS to drive this important developmental process.

## MATERIALS AND METHODS

### Growth conditions and root hair quantification

All Arabidopsis mutants were in the Col-0 background. The *etr1-7, etr1-3* (AT1G66340), *ein3-1* (AT3G20770) *eil1-1* (AT2G27050) double and *rhd2-6* (AT5G51060) mutants have all been described previously (Binder et al., 2007; Gayomba and Muday, 2020; Harkey et al., 2018; Hua and Meyerowitz, 1998). The ZAT12p-ROS construct was generously provided by Won-Gyu Choi (Lim et al., 2019). *prx44-2* (AT4G06010) and *prx73-4* (AT5G67400) were obtained from the Arabidopsis SALK center (SALK_057222C and SALK_020724, respectively), the mutations were verified by PCR, and homozygous lines were isolated. The transcriptional reporter *pPRX44*::GFP transgenic line was described previously (Marzol, 2021). Seeds were sterilized in 750% ethanol for approximately 5 minutes and grown on 100 × 15 mm Petri dishes containing 25 mL media. Seedlings were grown on 1x MS supplemented with 1% sucrose, vitamins (1 µg/ml thiamine, 0.5 µg/ml pyridoxine and 0.5 µg/ml nicotinic acid), 0.05% MES (w/v) and 0.8% (w/v) agar. Media pH was 5.5. Micropore tape was used to seal the top of the Petri dish and plated seeds were stratified at 4° C in darkness for 2 days. Plants were grown under 24h light from T5 fluorescent lights at 120-150 μmol photons m^-2^ s^-1^.

### ACC and ethylene treatments

ACC stocks at 200 mM were prepared using ACC hydrochloride diluted in H_2_O. ACC was then added directly to media to yield of final concentration of 0.7 µM. Seeds were grown on control plates as described above for 5 days and then transferred to plates containing either 0.7 μM of ACC or transferred to new plates and placed in an air tight tank containing 0.5 ppm ethylene gas. Seedlings were then left under the light conditions described above for 4 hours and then used for experiments.

### Quantification of root hair number and root hair length

Seedlings were grown for 5 days on control media and then transferred to media containing 0.7 µM ACC and grown for 4 hours or other indicated times. To examine root hairs, 5-day old seedlings were imaged using bright-field on an Axio Zoom V16 stereomicroscope. Extended depth of focus was used to combine z-stack images. Root hairs that were at stage +1 or +2 (or later stages) as defined by (Denninger et al., 2019) were quantified using Fiji/ImageJ software in three zones (0-500 μm, 500-1000 μm, 1000-1500 μm) starting from the root tip.

We also determined root hair length in the presence and absence of ACC treatment in Col-0 ad mutants. All root hairs in zone 2 that had begun to elongate (were at stages above +2) were measured using Fiji/ImageJ software. The length of each root in microns was determined. To obtain the histogram of distribution, the roots were separated into bins of different lengths, and the % of root hairs in each length bin was determined by dividing by the total number of root hairs in Zone 2 in that root.

### Confocal imaging of dyes and reporters of ROS levels

H_2_O_2_ was visualized with Peroxy-Orange 1 (PO1). PO1 was dissolved in DMSO to make a 500 μM stock and was further diluted in H_2_O to make a 50 μM working solution. Seedlings were incubated in PO1 for 15 minutes in the dark and were then rinsed with H_2_O and mounted in H_2_O for imaging. Control, ACC-treated and ethylene-treated seedlings were imaged on a Zeiss 880 laser scanning confocal microscope using a 10x objective. PO1 was excited with a 488 nm laser at 0.25% power and emission was collected between 544-695 nm. Images were analyzed using Fiji/ImageJ software. Plot profiles were taken across the epidermal cell files of maximum intensity projections using a 20-pixel line and values were averaged within each cell file. All images were captured at levels below saturation, although images shown in the manuscript were uniformly adjusted for brightness and contrast for clarity.

5-day old seedlings harboring the ZAT12p-ROS construct were mounted in H_2_O and excited with 488 and 561 nm lasers at 6% and 1.2% laser power, respectively. GFP emission was collected at 521 nm and mCherry emission was collected at 593 nm. Fiji/ImageJ software was used to generate two single channel images to form individual GFP and mCherry channels. Plot profiles were taken using a 250-pixel wide line to measure fluorescence of the entire root. Measurements were taken starting at 200 μm back from the root tip and ending at 1500 μm. Ratios were generated by dividing the GFP channel by mCherry using the Image Calculator tool (Process/Image Calculator).

### PI staining and *proPRX44*::GFP imaging

Cell walls were stained with 0.5 µg/mL propidium iodide (PI) dissolved in H_2_O. 5 day old seedlings treated with and without ACC were incubated in PI for 4 minutes before imaging. Fluorescence was visualized using a 561 nm laser at 0.15% and emission spectra set to 561-695 nm. These settings were used for all images. Optical slices of the top section of the root to show epidermal cells and maximum intensity projections of z-stack images are shown in Fig. S1. Fiji software was used to measure cell length of 3 epidermal cells per root using optical slices of the top section of the root. Cells were chosen at the bottom of zone 2 (cell 1, in the middle (cell 2), and at the top (cell 3). 4 cells of each of the 3 cell types were averaged and are shown in FS1. 6-8 roots of both control and ACC treated seedlings. Transgenic seeds harboring *pPRX44*::GFP were obtained from the lab of Dr. Jose Estevez, 5-day old seedlings were mounted in H_2_O and excited with a 488 nm laser at 5% and emission was collected between 490-606 nm. Seedlings were then stained with PI.

### DNA extraction and PCR for mutant genotype analysis

T-DNA insertion lines were grown for approximately 2 weeks on media described above, then transferred to soil and grown under 24h light at 50-80 μmol photons m^-2^ s^-1^. Leaves were harvested from 6 week old plants and stored in eppendorf tubes at −20^°^ C prior to DNA extraction. Frozen leaves were ground in DNA extraction buffer (100 mM Tris-HCl pH 8.0, 10 mM Na2EDTA, 100 mM LiCl_2_, 1% (w/v) SDS). Samples were then washed once with isopropanol and three times with 80% (v/v) ethanol. Finally, DNA pellets were dried, resuspended in sterile H_2_O and stored at −20^°^ C. Each PCR reaction contained 1X GoTaq Polymerase, 1 µM of each primer, 1 uL of DNA and 5 µL H_2_O. To confirm that mutants were homozygous for the desired T-DNA insertion, one reaction was performed with primers (left primer and right primer) flanking the left and right sides of in-tact genes while one reaction was performed with the right primer and LBb1.3, which is a primer specific for the left border of the T-DNA insertion. PCR products were run on 0.8 % (w/v) agarose gels containing 0.002% SERVA DNA stain G.

### RNA Isolation and Quantitative Real Time-PCR

RNA was isolated from seedlings grown on a nylon filter as described previously (Harkey et al., 2018). 5-days after germination seedlings were transferred to either control media or media containing 0.7 μM ACC and were grown for 4 hours. Roots were then cut and flash frozen in liquid nitrogen then ground. RNA was isolated using the Qiagen plant RNeasy kit protocol and RNA was DNase treated.

cDNA synthesis was performed using the RevertAid RT Reverse Transcription kit (ThermoFisher). qRT-PCR analysis using this cDNA was performed on a QuantStudio real time PCR machine using SYBR green detection chemistry. Primers specific to *PRX44* (FP: 5’-CAA GAG ACT CGG TCG CAT TAG-3’ and RP: 5’-TTG TTG GTC CGG GTA AGT TC-3’) and *RBOHC* (FP: 5’-CAA GGA ACA AGC CCA ACT AAA-3’ and RP: 5’-TTC TAT TGG GTT ACG CGT GAG-3’) were used and relative transcript abundance was quantified using ACT2 primers using comparative Ct analysis (ΔΔCt) to determine the relative quantity of target transcripts.

### NADPH oxidase/RBOH enzyme assays

Protein extract was isolated from seedlings grown on a nylon filter as described previously (Harkey et al., 2018). After 2 days of stratification and 7 days of growth under conditions described above, the nylon was transferred to growth medium with and without 0.7 μM ACC for 4 hours. Roots were then cut from seedlings and flash frozen in liquid nitrogen. Frozen samples were ground in liquid nitrogen using a mortar and pestle. RBOH extraction buffer (50 mM Tris-HCl pH 7.5, 0.1 mM EDTA, 0.1% (v/v) Triton X-100, 1 mM MgCl_2_, 10% (v/v) glycerol) was then added to a plant material/buffer ratio of 1:3 (w/v). Samples were centrifuged and supernatant was collected and desalted and concentrated using the Amicon Ultra-0.5 Centrifugal Filter devices. RBOH reaction mixture (50 mM Tris-HCl pH 7.5, 1 mM CaCl_2_, 0.1 mM nitroblue tetrazolium (NBT), 0.1 mM NADPH) was then mixed with protein extract at a 1:1 ratio. The reduction of NBT to monoformazan was monitored spectrophotometrically at 530 nm. Monoformazan concentrations were calculated using a 12.8 mM^-1^cm^-1^ extinction coefficient. This assay was adapted in sweet peppers and has previously been described (Chu-Puga et al., 2019).

## ACKNOWLEDGMENTS

We would like to thank Dr. Brad Binder (University of Tennessee) and all Muday lab members for their editorial input, and Dr. Glen Marrs and Dr. Heather Brown-Harding (Wake Forest University) for their help with microscopy.

## COMPETING INTERESTS

The authors have no competing interests.

### FUNDING

This project was supported by the National Science Foundation (MCB-1716279 to GKM) and a USDA Predoctoral Fellowship (NIFA 2021-67034-35113 to REM). Dr. Jose Estevez and Dr. Eliana Marzol are investigators of the National Research Council (CONICET) from Argentina supported by grants from ANPCyT (PICT2019-0015 to J.M.E. and PICT2018-0577 to E.M.) and ANID – Programa Iniciativa Científica Milenio ICN17_022, NCN2021_010 and Fondo Nacional de Desarrollo Científico y Tecnológico [1200010] to J.M.E.

### DATA AVAILABILITY

All data in this manuscript are included in the text or in the supplementary file.

