## Supplemental Table and Figures for "Ethylene signaling increases reactive oxygen species accumulation to drive root hair initiation in Arabidopsis"

Table S1. Transcript abundance changes in genes encoding RBOHC, class III peroxidases, and root hair patterning genes

| | | Signal log ratio values after ACC treatment (hours)[a] | | | | | | | |
|---|---|---|---|---|---|---|---|---|---|
| Gene Symbol | Locus ID | 0 | 0.5 | 1 | 2 | 4 | 8 | 12 | 24 |
| RHD2 (RBOHC)[b] | AT5G51060 | -0.09 | 0.32 | 0.28 | 0.33 | 0.37 | 0.05 | -0.06 | 0.16 |
| PRX44[c] | AT4G26010 | 0.40 | -0.78 | 0.47 | 0.67 | 0.68 | 0.84 | 0.89 | 1.19 |
| PRX69[c] | AT5G64100 | -0.09 | 0.51 | 0.80 | 0.96 | 1.23 | 1.32 | 1.50 | 0.80 |
| PRX73[b] | AT5G67400 | 0.6 | -1.1 | 0.3 | 0.39 | 1.42 | 1.2 | 1.2 | 1.43 |
| PRX39[c] | AT4G11290 | 0.07 | 0.00 | 0.15 | 0.24 | 0.86 | 0.97 | 1.51 | 0.62 |
| PRX25[c] | AT2G41480 | -0.78 | 0.60 | 0.20 | -0.71 | 0.28 | -0.20 | -0.08 | 0.53 |
| PRX72[c] | AT5G66390 | -0.09 | 0.11 | -0.05 | -0.12 | -0.43 | -0.94 | -0.96 | -1.02 |
| PRX64[c] | AT5G42180 | 0.08 | -0.35 | 0.02 | 0.00 | -0.44 | -0.81 | -0.84 | -1.06 |
| PRX66[c] | AT5G51890 | 0.01 | -0.02 | 0.00 | 0.21 | -0.27 | -0.45 | -0.82 | -1.32 |
| ROP2[b] | AT1G20090 | 0.08 | -0.04 | -0.13 | -0.11 | -0.12 | -0.31 | -0.22 | -0.28 |
| ROPGEF3[b] | AT4G00460 | 0.03 | 0.00 | 0.28 | -0.08 | -0.06 | -0.11 | -0.12 | -0.12 |
| ROPGEF4[b] | AT2G45890 | 0.26 | -0.59 | -0.08 | 0.21 | -0.04 | -0.22 | -0.30 | -0.34 |
| GL2[d] | AT1G79840 | -0.06 | -0.46 | -0.36 | -0.23 | -0.42 | -0.32 | -0.44 | -0.29 |
| RHD6[b] | AT1G66470 | 0.10 | -0.56 | 0.06 | -0.01 | -0.17 | -0.31 | -0.24 | -0.09 |
| RSL4[b] | AT1G27740 | -0.25 | 0.00 | -0.19 | -0.10 | 0.04 | -0.01 | -0.30 | -0.22 |

[a]Signal log ratio (SLR; log base 2 of fold change) was calculated for each transcript relative to an averaged time matched control (Harkey et al 2018); [b]transcripts that did not change significantly in response to ACC in either the DE or PA filtering; [c]transcripts that were differentially regulated in response to ACC; [d]transcripts that were present (above background) at some time points while absent at others (PA).

Signal log ratio

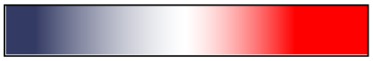

-1.5 0 1.5

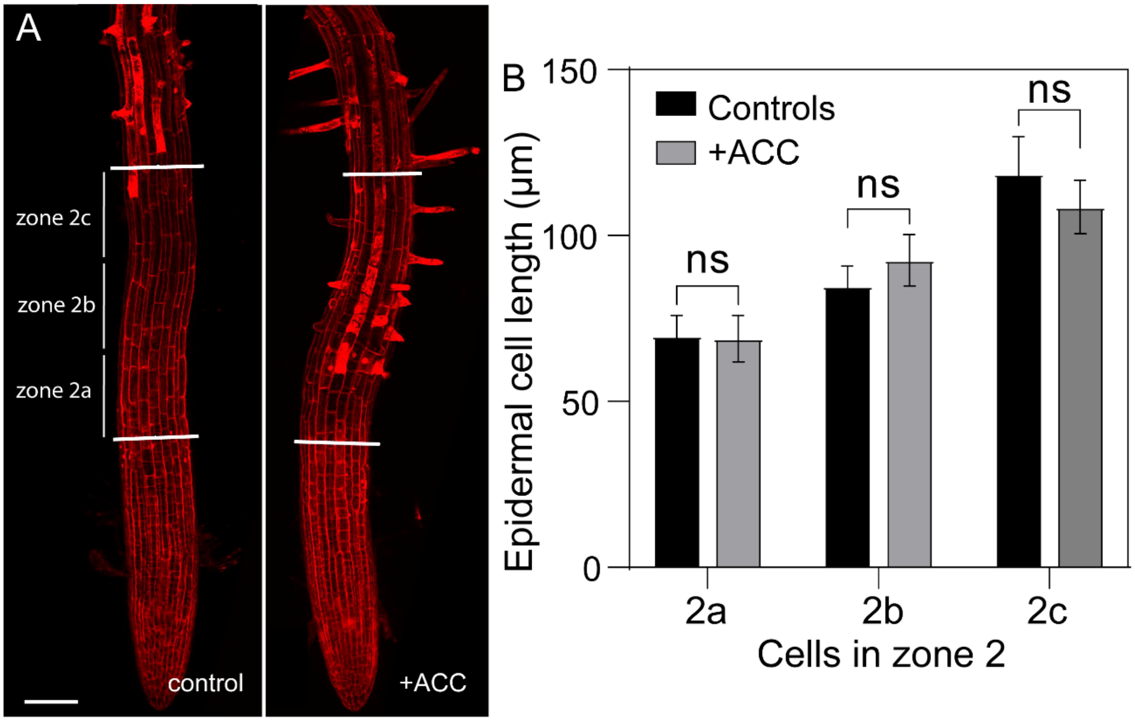


**Figure S1. Epidermal cell length did not change after 4 hours of ACC treatment** (A) Representative images of Col-0 seedlings treated with and without ACC for 4 hours and stained with propidium iodide (PI). Scale bar is 100 µm. (B) Quantification of an average length of 12-18 epidermal cells. 3 cells per zone were analyzed, a cell towards the bottom of the zone 2 (cell 2a), a cell in the middle (cell 2b) and a cell at the top of the zone (cell 2c). Data are ± SEM of 3 independent experiments (n=12-15 seedlings per experiment). Data was statistically analyzed using two-way ANOVA followed by Tukey's multiple comparisons test.

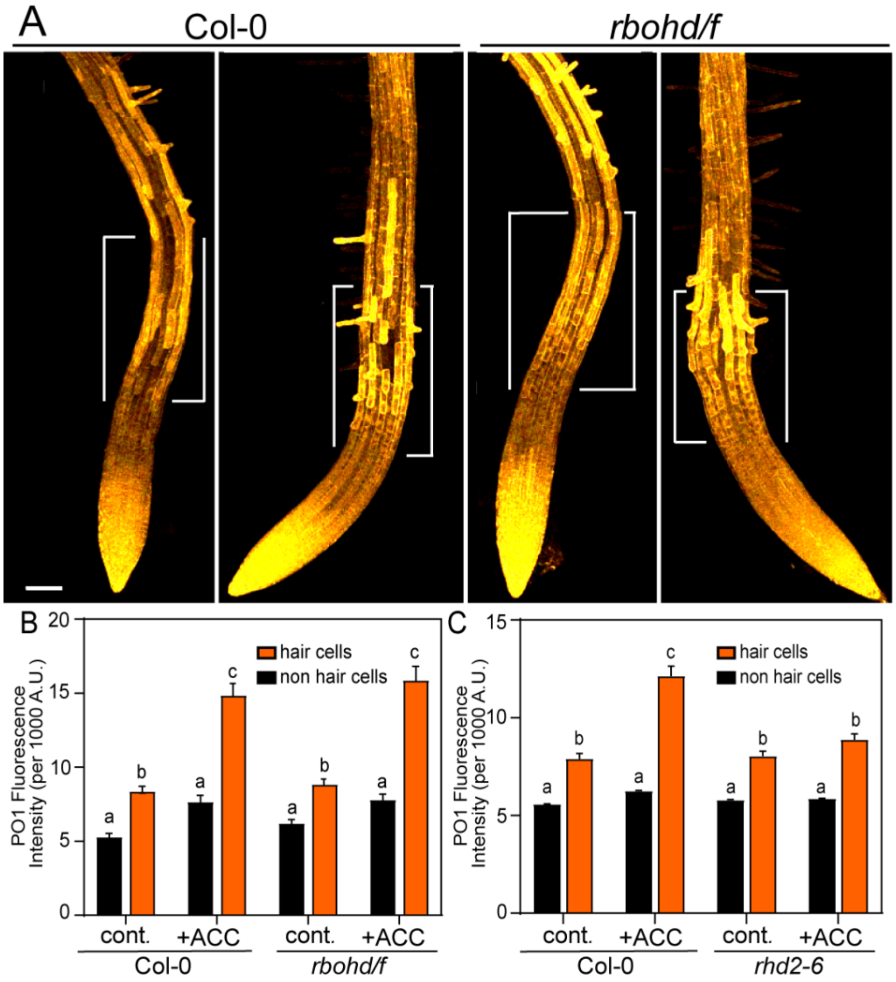


**Figure S2. RBOHD and RBOHF do not contribute to ethylene-induced ROS synthesis in root hair cells** (A) Representative images of PO1 epidermal fluorescence in Col-0 and *rbohd/f* treated with and without ACC for 4 hours. Scale bar is 100 µm. (B) Quantification of PO1 fluorescence intensity in hair cells (1, 3, and 5) and non-hair cells (2, 4) of Col-0 and *rbohd/f* treated with and without ACC. Data are means ± SEM. of 3 experiments (n=12-18 seedlings/experiment). Columns with different letters indicate statistical significance compared to other hair cells as determined by two-way ANOVA followed by Tukey's multiple comparisons test. (C) For comparison, quantification of PO1 fluorescence intensity in hair cells and non-hair cells of Col-0 *rhd2-6 (rbohc)* treated with and without ACC from Figure 6E is shown. Data are means ± SEM. of 3 experiments (n=12-18 seedlings/experiment). Columns with different letters indicate statistical significance compared to other hair cells as determined by two-way ANOVA followed by Tukey's multiple comparisons test.

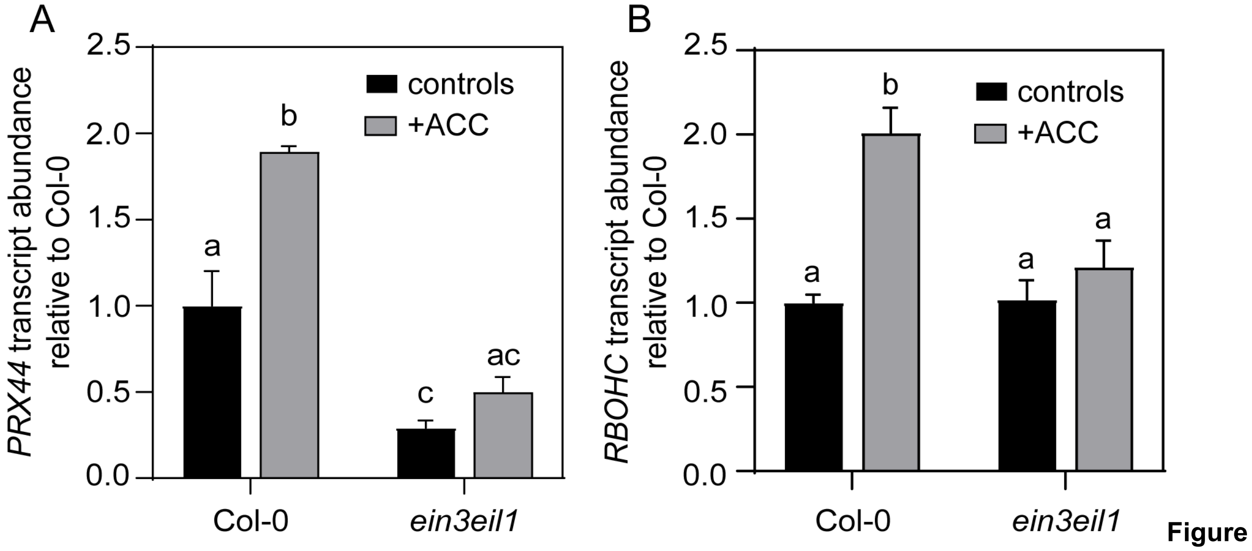


**Figure S3. *PRX44* and *RBOHC* transcript abundance is regulated by EIN3/EIL1 (A)** *PRX44* and **(B)** *RBOHC* transcript abundance in Col-0 and *ein3eil1* mutants treated with and without 0.7 µM ACC for 4 hours as determined by qRT-PCR. Averages of 3 replicates are reported. Different letters represent statistical significant differences as determined by two-way ANOVA followed by Tukey's multiple comparisons test.

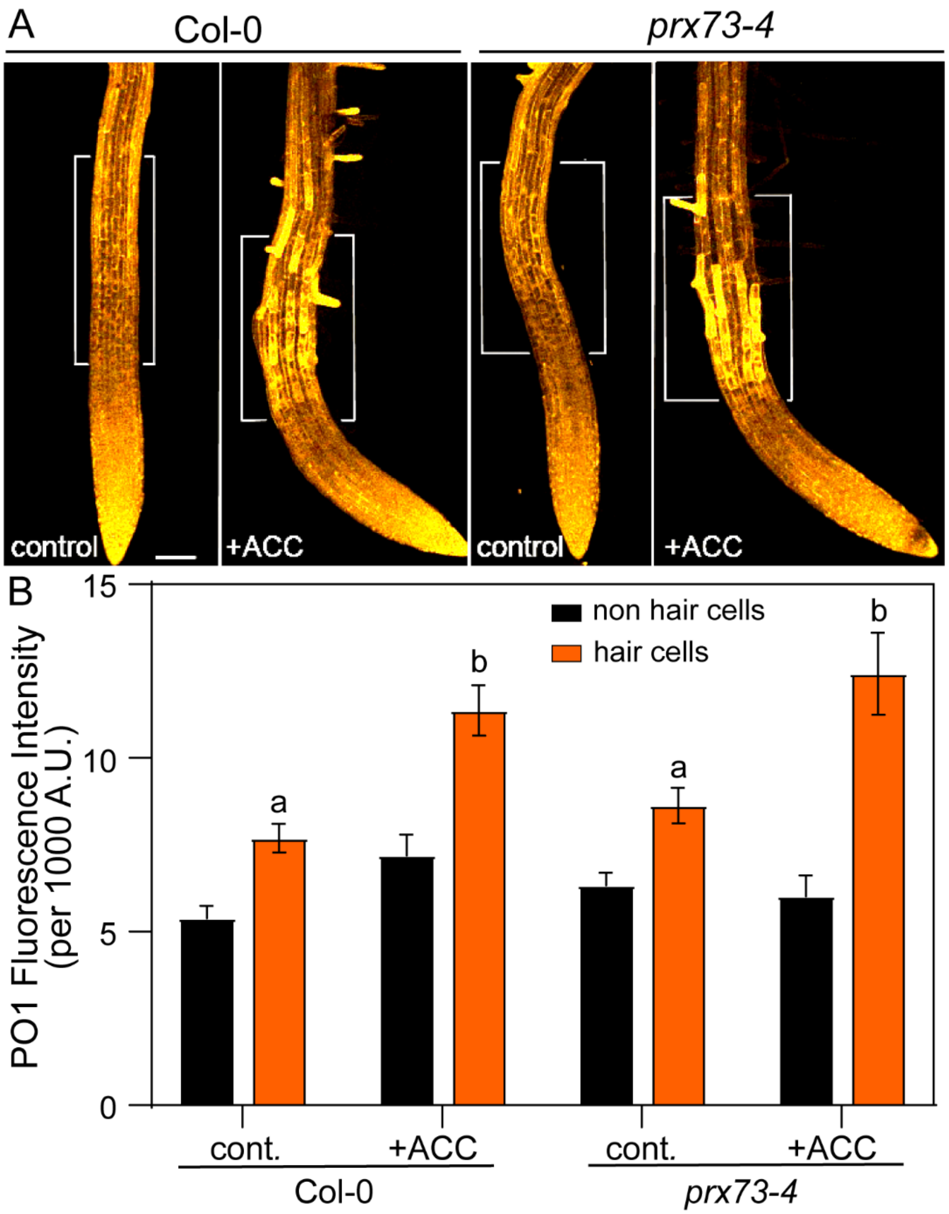


**Figure S4. PRX73 does not contribute to ACC-induced ROS synthesis in root hair cells** (A) Representative images of PO1 epidermal fluorescence in Col-0 and *prx73-4* treated with and without ACC for 4 hours. Scale bar is 100 μm. (B) Quantification of PO1 fluorescence intensity in hair cells (1, 3, and 5) and non-hair cells. Data are means ± SEM. of 3 independent experiments (n=15-18 seedlings per experiment). Columns with different letters indicate statistical significance compared to other hair cells as determined by two-way ANOVA followed by Tukey’s multiple comparisons test.
